# Beyond financial conflicts of interest: Institutional oversight of faculty consulting agreements at schools of medicine and public health

**DOI:** 10.1101/394585

**Authors:** Michelle M. Mello, Lindsey Murtagh, Steven Joffe, Patrick L. Taylor, Yelena Greenberg, Eric G. Campbell

## Abstract

**Importance:** Approximately one-third of U.S. life sciences faculty engage in industry consulting. Despite reports that consulting contracts often impinge on faculty and university interests, institutional approaches to regulating consulting agreements are largely unknown.

**Objective:** To investigate the nature of institutional oversight of faculty consulting contracts at U.S. schools of medicine and public health.

**Design:** Structured telephone interviews with institutional administrators. Questions included the nature of oversight for faculty consulting agreements, if any, and views about consulting as a private versus institutional matter. Interviews were analyzed using a structured coding scheme.

**Setting:** All accredited schools of medicine and public health in the U.S.

**Participants:** Administrators responsible for faculty affairs were identified via internet searches and telephone and email follow-up. The 118 administrators interviewed represented 73% of U.S. schools of medicine and public health, and 75% of those invited to participate.

**Intervention:** Structured, 15-30 minute telephone interviews.

**Main outcomes and measures:** Prevalence and type of institutional oversight; responses to concerning provisions in consulting agreements; perceptions of institutional oversight.

**Results:** One third of institutions (36%) required faculty to submit at least some agreements for institutional review and 36% reviewed contracts upon request, while 35% refused to review contracts. Among institutions with review, there was wide variation the issues covered. The most common topic was intellectual property rights (64%), while only 23% looked at publication rights and 19% for inappropriately broad confidentiality provisions. Six in ten administrators reported they had no power to prevent faculty from signing consulting agreements. Although most respondents identified institutional risks from consulting relationships, many maintained that consulting agreements are “private.”

**Conclusions and relevance:** Oversight of faculty consulting agreements at U.S. schools of medicine and public health is inconsistent across institutions and usually not robust. The interests at stake suggest the need for stronger oversight.

## Introduction

Approximately one-third of life sciences faculty engage in industry consulting [1], providing paid advice or services to companies whose activities relate to their areas of expertise.[1,2,3,4] Consulting activities can be valuable in advancing science and technology in medicine and the life sciences,[5] yet they create controversy because they may influence the conduct and reporting of research and undercut openness in science.[6,7,8,9,10] To date, the public conversation and resultant policy action have focused on financial conflicts of interest (fCOI). The potential for financial incentives to influence faculty to act in ways that are inconsistent with their duties to universities and research participants and contrary to the core values of science has led to a broad net of public and private oversight.[6,11,12]

Financial conflicts stemming from industry relationships, however, are not the only reason for concern. Both industry-sponsored research and private consulting relationships rely upon contracts between companies and faculty or their institutions that create legally enforceable obligations and rights. As with sponsored research,[13] companies might use consulting contracts to exert inappropriate influence over academic research and investigators.[14,15,16] For example, consulting contracts may require the company’s approval for the consultant to publish, even for work beyond the scope of the consultancy; restrict the consultant’s ability to make public statements or engage in projects that are inimical to the company’s interests; or give the company ownership of intellectual property generated during the period of the consultancy even if it arises from the consultant’s academic work.[17]

Although medical school administrators and attorneys report that consulting agreements often contain language that restricts faculty members’ academic freedom and may threaten the integrity of their research [18], institutions’ approaches to addressing such problems have rarely been systematically studied.[19] Available guidelines are limited and no regulatory statements address universities’ roles in managing nonfinancial aspects of consulting relationships. An Institute of Medicine committee and the Institute on Medicine as a Profession support institutional review of consulting contracts, but they offer no details concerning the nature of the review.[6,20] The Association of American Medical Colleges provides a list of “topics and questions to consider” that is “neither exhaustive nor exemplary.”[21] The American Association of University Professors simply advises that faculty should not sign consulting contracts that undercut their ability to express their opinions [11]; and guidelines from the Pew Charitable Trusts merely state that consulting contracts should have “clear deliverables” and compensation set at fair market value.[22] Responsibility for executing appropriate consulting agreements is largely devolved to individual faculty or supervisors, who may be unaware of the potentially significant legal implications of what they sign. Here, we report the first empirical findings concerning the extent to which U.S. schools of medicine and public health regulate the content of faculty consulting agreements.

## Materials and methods

### Sample

We interviewed administrators at accredited U.S. medical schools and schools of public health. To recruit respondents, we searched schools’ websites to identify individuals who, given their positions, were likely to be knowledgeable about faculty consulting. We requested an interview or referral to a more knowledgeable administrator at the same institution. Where persons initially contacted did not respond or declined participation without indicating whether they were an appropriate respondent, we identified another knowledgeable person at the school using information on the school’s website. Participants received a $20 incentive.

Oversight of consulting was sometimes centralized rather than managed separately within the medical and public health schools. For these “affiliated” schools, we interviewed one informant from the office conducting centralized oversight unless he/she indicated we should also speak to someone else. In calculating response rate, we counted affiliated schools as one institution, resulting in a denominator of 157 eligible persons (details in Appendix).

## Interviews

We conducted 15- to 30-minute telephone interviews in 2011 using a computer-assisted interview guide on the REDCap Survey platform. [23] Questions were developed based on a checklist of restrictive provisions developed by a major academic center and a past survey concerning sponsored research agreements. [13] Interviewers provided a definition of “consulting relationship” and distinguished it from sponsored research.

Interviews were conducted by one of three investigators, following training that included listening in on several interviews to achieve consistency in style. Interviewers took detailed notes in REDCap during the interview.

## Analysis

A detailed coding guide for free-text interview responses was created based on two investigators’ review of a sample of six schools’ interview notes and recollections of responses from other interviews. Each investigator generated a coding scheme independently and differences were discussed and resolved. The final coding guide was programmed into REDCap and each set of interview notes was coded by one of two investigators. The resulting quantitative data were analyzed using Stata 10 (College Station, TX). Multivariable logistic regression was used to examine school characteristics as predictors of oversight approach, applying a significance level of 0.05 in two-tailed tests. Some free-text responses were qualitatively analyzed. The study was approved by the Harvard School of Public Health institutional review board. All participants gave written informed consent to research participation.

## Results

### Sample characteristics

Interviews were completed with administrators representing 127 of 173 medical schools and schools of public health in the U.S. (73%) (Table 1). Of 157 eligible administrators, 118 (75%) participated. The most common job title was some variant of associate dean for research, but directors of offices of sponsored programs, research compliance and general counsel were also highly represented.

**Table 1.** Characteristics of institutions represented in key informant interview sample

|  |  |  |
| --- | --- | --- |
|  | No. | %<sup>a</sup> |
| Institutions represented | 127 | -- |
| Schools of medicine | 95 | 75% |
| Schools of public health | 32 | 25% |
| Number of administrators interviewed<sup>b</sup> | 118 | -- |
| Mean number per institution | 1.1 | -- |
| Schools' NIH funding rank |  |  |
| Schools of medicine: |  |  |
| Top 10% | 11 | 12% |
| 11%-51% | 42 | 44% |
| Bottom 50% | 40 | 42% |
| Not available | 2 | 2% |
| Schools of public health: |  |  |
| Top 10% | 4 | 13% |
| 11%-51% | 14 | 44% |
| Bottom 50% | 12 | 38% |
| Not available | 2 | 6% |

<sup>a</sup> Percentages may not sum to 100 due to rounding.
<sup>b</sup> At institutions' request, two informants were interviewed at each of 12 institutions. Fourteen administrators each had responsibility for oversight at two or more affiliated schools.

Fourteen key informants represented more than one school within their university. At 12 institutions, we interviewed two informants because administrators suggested we speak with someone at both the school and the university/health campus level. Their responses were merged because institutions were the unit of analysis.

### Prevalence and types of oversight approaches

About one third of institutions (36%) required faculty to submit consulting agreements for institutional review prior to execution; however, only about half of these (23 institutions) required review for all agreements (Table 2). The other 17 required review only if certain triggering conditions were present—for instance, the consulting activity was related to the faculty member’s research, or the faculty member opted to make the institution a party to the contract. At a third of institutions (36%), administrators would review faculty members’ consulting agreements upon request but did not require review. Thirty-nine institutions (35%) did not review consulting agreements even if asked. In multivariable logistic regression models controlling for NIH funding rank tercile and school type (medical versus public health), neither characteristic significantly predicted the likelihood of taking each approach to reviewing consulting agreements (mandatory, optional, or no review) (results not shown).

**Table 2.** Prevalence of institutional oversight approaches for faculty consulting agreements among schools of medicine and public health ^a^

| Type of oversight | No. | % |
| --- | --- | --- |
| **Mandatory review** | **40** | **36%** |
| All agreements reviewed | 23 | 21% |
| Under some circumstances | 17 | 15% |
| **Optional review available** | **40** | **36%** |
| When faculty member asks, but done purely as a favor | 38 | 34% |
| Under some conditions only | 3 | 3% |
| **No review available** | **39** | **35%** |
| **Other approaches** | **55** | **49%** |
| May be included in conflict-of-interest disclosure process | 22 | 20% |
| School tries to convert project to sponsored research; only reviews if converted | 13 | 12% |
| Addendum provisions required to be included | 7 | 6% |
| Addendum available listing recommended provisions | 7 | 6% |
| Other | 5 | 5% |

<sup>a</sup> Denominator for proportions (112) is the number of “affiliated schools” (universities where a single administrator handled matters for 2 or more schools) plus the number of “unaffiliated” schools of medicine plus the number of “unaffiliated” schools of public health. Percentages may not sum to 100 due to rounding or because response categories were not mutually exclusive (e.g., 7 schools coupled mandatory review for some types of agreements with optional review for others).

Many institutions described oversight approaches other than reviewing consulting agreements. Twenty-two (20%) said that information about restrictive provisions might be elicited during the school’s fCOI disclosure process, but acknowledged that this typically occurred after contract execution. Thirteen (12%) attempted to persuade faculty to convert consulting contracts to sponsored research agreements, which would be reviewed by the school’s sponsored programs office. Fourteen required or recommended that faculty attach a standard addendum to their consulting contracts containing generic provisions designed to protect the university’s and/or faculty member’s interests. Twenty-six institutions (23%) reported that they had no oversight mechanisms relating to restrictive provisions in consulting agreements, though they did have conflict-of-commitment policies.

### Qualifications of contract reviewers

The 73 institutions that reviewed consulting agreements on either an optional or a mandatory basis reposed responsibility for such review in a variety of types of administrators. Most common was the office of legal counsel (51%), followed by offices of research administration or industry relations (41%) and offices of technology transfer or intellectual property (30%). Smaller proportions used department chairs (10%), representatives from offices of the dean or president (12%), research compliance officers (12%), or fCOI committee staff (14%). Half (51%) required that reviewers have legal or risk-management training.

### Issues covered by institutional review

Among the 73 institutions that reviewed consulting agreements, the issues addressed by review varied widely (Table 3). The most common focus was protection of the university’s intellectual property rights (64%), followed by fCOI (29%), conflicts of commitment (29%), and whether services were being offered for fair market value (25%). Few institutions (7%) reviewed agreements to verify that required addenda had been attached, that the consulting arrangement did not violate applicable law (16%), or that the arrangement would not adversely affect trainees (4%).

**Table 3.** Issues covered in institutional review of consulting agreements (*n*=73) ^a^

| Issues included in review | No. | % |
| --- | --- | --- |
| **Predominantly institutional interests** |  |  |
| Intellectual property rights | 47 | 64% |
| Use of institution's property | 11 | 15% |
| Use of institution's name in consulting activity | 8 | 11% |
| Institution is not a party to the agreement | 7 | 10% |
| Effect on students/teaching | 3 | 4% |
| Inappropriate disclosure of information owned by institution | 2 | 3% |
| **Conflicts between institution's and faculty member's interests** |  |  |
| Potential conflicts of interest | 21 | 29% |
| Conflicts of commitment | 21 | 29% |
| Compliance with policies on consulting/outside activities | 14 | 19% |
| Whether proposed activity impermissibly overlaps with faculty member's institutional work/role | 16 | 22% |
| Existence of statement that obligations to school take precedence over obligations to company | 11 | 15% |
| Whether proposed activity is consistent with institution's mission | 2 | 3% |
| **Predominantly faculty member's interests** |  |  |
| Publication restrictions | 17 | 23% |
| Liability issues | 16 | 22% |
| Confidentiality of information received through the consulting work | 14 | 19% |
| Noncompete clauses affecting faculty member's future research activities | 10 | 14% |
| Choice of law / dispute resolution provisions | 5 | 7% |
| Issues raised by faculty member as concerning | 2 | 3% |
| **Other issues** |  |  |
| Whether services are provided for fair market value | 18 | 25% |
| Violation of state or federal laws/policies (e.g., NIH policy) | 12 | 16% |
| General appropriateness of consulting arrangement | 9 | 12% |
| Whether faculty member is asked to endorse a product | 6 | 8% |
| Addendum or other required provisions are included | 5 | 7% |
| Termination provisions | 3 | 4% |
| Unclear from interview responses | 9 | 12% |

<sup>a</sup> Denominator for proportions (73) is the number of schools that conducted some type of mandatory or voluntary review. Percentages may not sum to 100 due to rounding or because response subcategories are not mutually exclusive. Table excludes some issues mentioned by only one respondent

Review rarely included matters that predominantly affected the faculty member’s interests, rather than the university’s. Strikingly, less than a quarter of institutions examined consulting contracts for restrictions on publication rights. About 22% looked for provisions that could expose faculty to liability risk. Less than a fifth looked at the scope of confidentiality provisions. Only 14% looked for noncompete clauses that could affect the faculty member’s future research activities. In general, the higher the administrative level at which review took place, the more inclusive was the range of issues covered by the review.

### Responses to problematic provisions in consulting agreements

When reviewers identified a seemingly problematic provision in a consulting contract, only some took assertive action (Table 4). Twenty-two of 73 institutions (30%) told the faculty member the provision must be changed and had the faculty member negotiate with the company, and 22% were willing to negotiate directly with the company. Many others referred the matter to legal counsel or senior university administrators for follow-up. Only 38% reported having the authority to prevent the faculty member from entering into the consulting relationship if their concerns were not resolved.

**Table 4.**
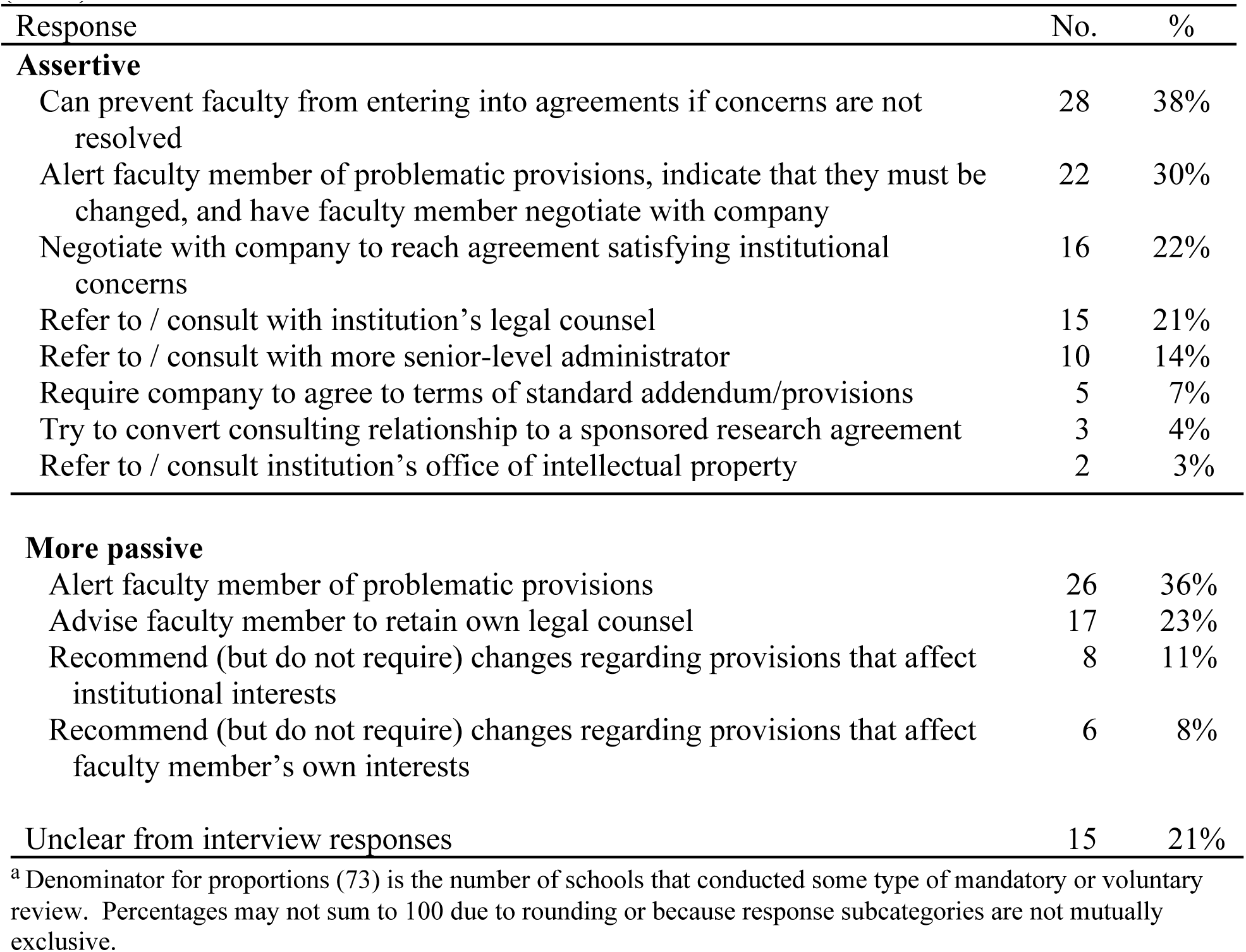
Institutional reviewers’ responses to troubling provisions in consulting agreements (*n*=73) ^a^

| Response | No. | % |
| --- | --- | --- |
| **Assertive** |  |  |
| Can prevent faculty from entering into agreements if concerns are not resolved | 28 | 38% |
| Alert faculty member of problematic provisions, indicate that they must be changed, and have faculty member negotiate with company | 22 | 30% |
| Negotiate with company to reach agreement satisfying institutional concerns | 16 | 22% |
| Refer to / consult with institution's legal counsel | 15 | 21% |
| Refer to / consult with more senior-level administrator | 10 | 14% |
| Require company to agree to terms of standard addendum/provisions | 5 | 7% |
| Try to convert consulting relationship to a sponsored research agreement | 3 | 4% |
| Refer to / consult institution's office of intellectual property | 2 | 3% |
| **More passive** |  |  |
| Alert faculty member of problematic provisions | 26 | 36% |
| Advise faculty member to retain own legal counsel | 17 | 23% |
| Recommend (but do not require) changes regarding provisions that affect institutional interests | 8 | 11% |
| Recommend (but do not require) changes regarding provisions that affect faculty member's own interests | 6 | 8% |
| Unclear from interview responses | 15 | 21% |

<sup>a</sup> Denominator for proportions (73) is the number of schools that conducted some type of mandatory or voluntary review. Percentages may not sum to 100 due to rounding or because response subcategories are not mutually exclusive.

Commonly, reviewers simply alerted faculty to problematic provisions and left the matter in the faculty member’s hands (36%). Seventeen institutions (23%) advised faculty to hire an outside attorney to resolve the issue.

### Perceptions of the need for institutional oversight

When asked to characterize how they perceived faculty consulting relationships to affect the institution’s interests, administrators identified both positive and negative effects. The most frequently mentioned benefits were helping to disseminate knowledge or speed research translation (35%), building external relationships (26%), raising the profile of the institution (21%) or faculty member (15%), giving faculty real-world experience (19%), creating research, educational, and funding opportunities (18%), and allowing faculty to supplement their income, which helped with retention (10%).

Most respondents (84%) recognized one or more potential negative implications of consulting relationships for the institution. The most common theme was that consulting relationships could restrict academic freedom, research activities, and/or research integrity (63%). Thirty-eight percent felt consulting could influence how faculty carry out their institutional roles and duties and 36% remarked that consulting activities could threaten the integrity of the institution or trust in its teaching or research. Similarly, many mentioned that consulting relationships could damage the institution’s reputation (30%), create conflicts of commitment (27%), or threaten the institution’s intellectual property rights (20%).

Institutions that required review of consulting contracts pointed to these risks when explaining the reasons for their approach (Table 5). Many expressed the desire to avoid legal problems or public scandals over faculty activities, while a few pointed to the need to safeguard the university’s intellectual property or voiced a sense that mandatory review of consulting contracts was the responsible thing to do.

**Table 5.**
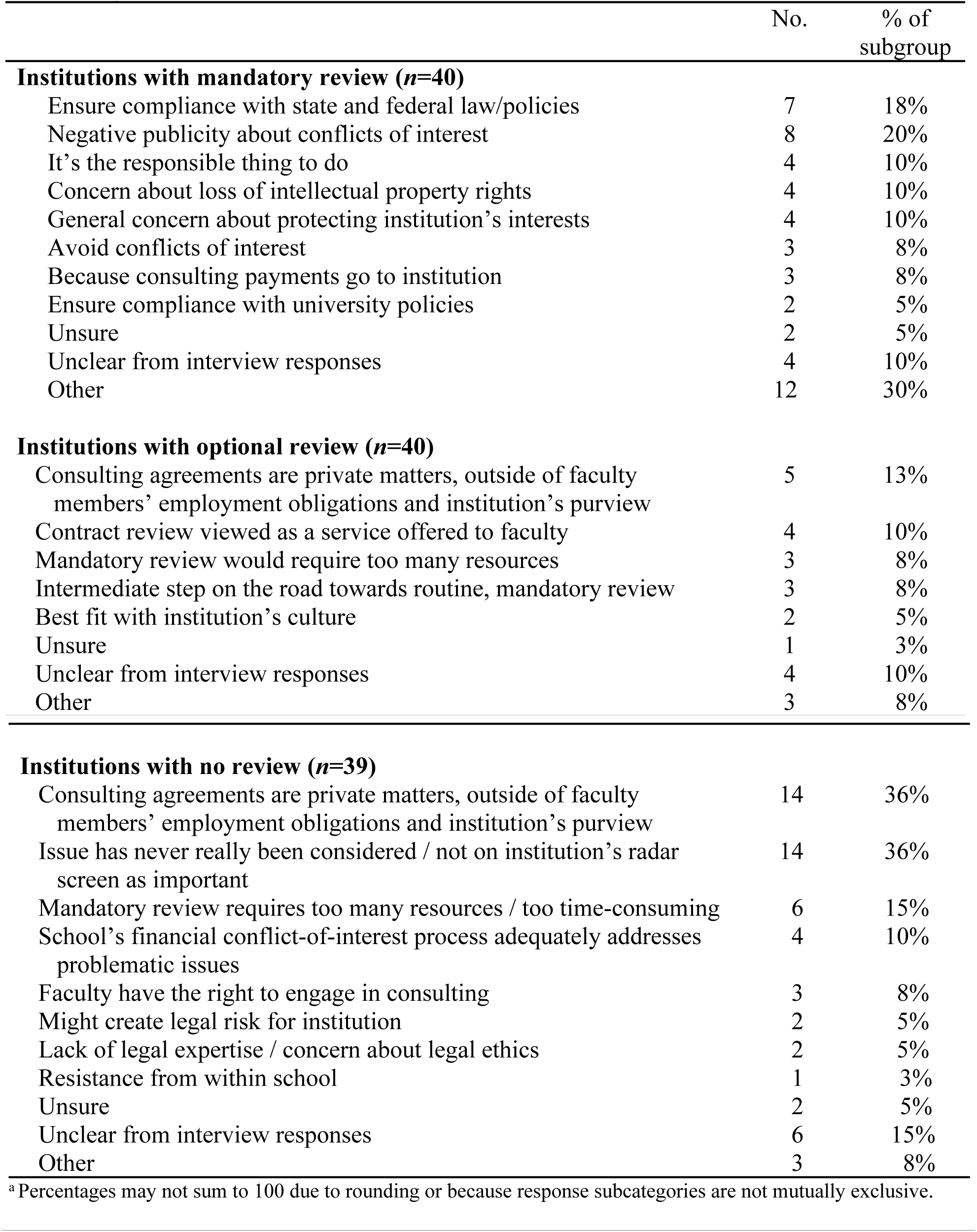
Institutions’ reasons for adopting particular approaches to review of consulting agreements^a^

|  | No. | % of subgroup |
| --- | --- | --- |
| **Institutions with mandatory review (n=40)** |  |  |
| Ensure compliance with state and federal law/policies | 7 | 18% |
| Negative publicity about conflicts of interest | 8 | 20% |
| It's the responsible thing to do | 4 | 10% |
| Concern about loss of intellectual property rights | 4 | 10% |
| General concern about protecting institution's interests | 4 | 10% |
| Avoid conflicts of interest | 3 | 8% |
| Because consulting payments go to institution | 3 | 8% |
| Ensure compliance with university policies | 2 | 5% |
| Unsure | 2 | 5% |
| Unclear from interview responses | 4 | 10% |
| Other | 12 | 30% |
| **Institutions with optional review (n=40)** |  |  |
| Consulting agreements are private matters, outside of faculty members' employment obligations and institution's purview | 5 | 13% |
| Contract review viewed as a service offered to faculty | 4 | 10% |
| Mandatory review would require too many resources | 3 | 8% |
| Intermediate step on the road towards routine, mandatory review | 3 | 8% |
| Best fit with institution's culture | 2 | 5% |
| Unsure | 1 | 3% |
| Unclear from interview responses | 4 | 10% |
| Other | 3 | 8% |
| **Institutions with no review (n=39)** |  |  |
| Consulting agreements are private matters, outside of faculty members' employment obligations and institution's purview | 14 | 36% |
| Issue has never really been considered / not on institution's radar screen as important | 14 | 36% |
| Mandatory review requires too many resources / too time-consuming | 6 | 15% |
| School's financial conflict-of-interest process adequately addresses problematic issues | 4 | 10% |
| Faculty have the right to engage in consulting | 3 | 8% |
| Might create legal risk for institution | 2 | 5% |
| Lack of legal expertise / concern about legal ethics | 2 | 5% |
| Resistance from within school | 1 | 3% |
| Unsure | 2 | 5% |
| Unclear from interview responses | 6 | 15% |
| Other | 3 | 8% |

<sup>a</sup> Percentages may not sum to 100 due to rounding or because response subcategories are not mutually exclusive.

A view that consulting agreements are private matters, outside of faculty members’ employment obligations and the university’s purview, was the primary reason that institutions made contract review optional (13%) or unavailable (36%). However, more than a third (36%) of the schools at which review was unavailable indicated that the issue of restrictive provisions in these contracts simply had not been on their radar screens. A minority of schools that did not provide review gave substantive reasons for rejecting that approach (Table 5)—for example, it would create a professional ethics problem for the university’s attorney, whose client was the institution, not individual faculty.

## Discussion

Universities and the public stand to lose when contractual relationships between faculty and companies are not carefully managed. Restrictive provisions in consulting agreements may jeopardize the progress of science by shifting intellectual property rights and restricting faculty members’ ability to publish scholarly work, engage in free intellectual discourse, pursue lines of scientific inquiry, and meet responsibilities to trainees.[15] Because consulting contracts create legally enforceable obligations that dictate behavior, not just incentives that may influence behavior, they are potentially of even greater concern than fCOI.

A lawsuit involving Stanford University illuminates the stakes.[17,24] The case arose after a research fellow employed by Stanford sojourned at a biotechnology company and subsequently developed an HIV testing method that built on his work during that time. His employment contract assigned his rights in inventions to Stanford. When Stanford sued to enforce its patents on the test, the company’s new owner responded that the researcher had signed a contract assigning the company his rights to inventions made during his time there. Resolving the conflicting contracts, the Supreme Court held in 2011 that the rights belonged to the company.

As this case demonstrates, the obligations that researchers assume in consulting agreements may cost universities dearly.[25] Moreover, the terms of consulting agreements may undercut the governance structures for collaborative research created by public and other funders, journal editors, and the law. They may, for instance, disrupt presumptions about authorship, intellectual property, and public disclosure obligations. Restrictive provisions in consulting agreements can also harm students and academic collaborators—for example, by signing away their rights in collaboratively developed inventions or imposing confidentiality obligations on them without their knowledge.

Previous research has explored institutional oversight of fCOI [6,22,26,27,28,29,30,31,32] and clinical trial agreements.[13,33,34,35,36] Our own work has examined normative beliefs about regulating consulting agreements among administrators at medical schools that have taken a particularly active approach.[19] The present study is the first, however, to systematically examine norms and practices relating to consulting oversight across U.S. medical schools and schools of public health.

### Shortcomings of current oversight

The important interests at stake call into question the traditional view of consulting agreements as private arrangements subject only to self-regulation by faculty and companies. In investigating whether practices among schools of medicine and public health reflect the traditional view, our study revealed several interesting findings.

First, there is heterogeneity in schools’ approaches to regulating the terms of consulting agreements. Schools are split between requiring institutional review of agreements, offering it as an option, and declining to provide review. Higher research intensity (NIH funding rank) did not predict approach. Rather, respondents attributed decisions to whether the potential risks of faculty consulting were on the institution’s “radar screen” and the extent to which institutional culture enshrined the view that consulting activities are private. In short, institutions lack a shared norm that they are justified in regulating this area at all, much less in a particular way.

Some institutions reported using other approaches instead of contract review, such as providing a standard addendum of provisions to be included in agreements. These mechanisms are weak compared to reviewing contracts, however. Providing an addendum does not ensure that faculty will include it, and beliefs that the fCOI disclosure process would identify restrictive contractual provisions seem misplaced in light of the rarity with which contracts are submitted. Even among schools that review contracts, there was substantial variation in what their review covered and how they responded to problematic provisions.

Second, contract review often focuses on protecting the institution’s own interests. In contrast to the two thirds of reviewing institutions that looked for provisions relating to intellectual property rights, less than a quarter looked for inappropriate restrictions on a faculty member’s ability to publish, provisions placing faculty at liability risk, or inappropriately broad confidentiality provisions. Review was frequently conducted by technology transfer offices, whose remit is to protect the university’s intellectual property. Such offices have little incentive to promote publication rights because publicizing inventions can undermine their patentability.

Third, many institutional administrators articulated conflicting views regarding whether universities should regulate this area. Many characterized consulting contracts as “private” and outside the institution’s purview, yet recognized that they can implicate the university’s interests in numerous, important ways. This dissonance may reflect more than reluctance to intrude into faculty members’ “private time,” which could affect schools’ ability to attract and retain top faculty. It may also spring from worries that reviewing consulting contracts could make the university vulnerable to lawsuits relating to those agreements.

Our study has limitations. Despite the high response rate, nonresponse bias may have affected our results. Interviews were conducted in 2011 and institutions subsequently may have changed their approaches, although we have no reason to think many have done so. Finally, interviews were not fully transcribed and nuances of responses could have been missed in notetaking.

### Strengthening oversight

Our findings suggest that oversight of faculty consulting agreements at most U.S. medical schools and schools of public health is highly variable and usually not robust. The evidence that consulting contracts often contain restrictive provisions and that such provisions can lead to harm is largely anecdotal [14,37,38], but the potential for harm and the spottiness of existing review practices raise questions about whether greater oversight should be exercised.

Management approaches could range from faculty training to mandatory review of consulting agreements.[19] Approaches that vest discretion in faculty to seek review may prove ineffectual because faculty may not appreciate the risks involved [16] even with educational outreach from the university, and have a countervailing financial interest in proceeding with the consulting relationship and avoiding the hassle of contract review. Faculty with the most problematic agreements may be the least willing to expose themselves to scrutiny.

One solution would be a “pay or play” policy in which universities would require faculty either to submit their consulting agreement for university review or attest that it was reviewed by a qualified attorney. The university could maintain a list of attorneys it has educated about its perspective on potentially problematic contractual provisions. The cost of external legal review could be built into faculty members’ consulting fees.

Requiring legal review of consulting contracts would likely meet with resistance from faculty, particularly if applied to consultancies with low remuneration. However, the history of fCOI regulation suggests this is no reason to abstain from oversight and that resistance would dissipate as institutions’ new role becomes culturally engrained. It also suggests that intervention from regulators and stronger guidance from professional organizations may be necessary to harmonize institutional approaches.

The irony of not regulating consulting contracts because they are “private” is that there is no obligation more fundamental for a tax-exempt organization than to be operated for the public benefit, and inappropriate contracts may divert institutional resources away from public purposes. Greater recognition of the ways in which faculty members’ putatively private consulting activities implicate public and institutional interests can promote the integrity of these valuable but ethically fraught relationships.

## Acknowledgments

The authors gratefully acknowledge Marc Lipsitch and David Korn for helpful comments on the project’s approach and findings and Aurora DeMattia for project assistance.

